# Correction of the Framingham Risk Score Data Reported in SPRINT

**DOI:** 10.1101/235358

**Authors:** Frederick Warner, Sanket S. Dhruva, Joseph S. Ross, Pranammya Dey, Karthik Murugiah, Harlan M. Krumholz

## Abstract

This report describes an error in the Framingham Risk Score data presented in the original SPRINT publication.^1^ The data, presented in Table 1 of the main SPRINT publication in the *New England Journal of Medicine* and made available to SPRINT Challenge participants, incorrectly calculated the level of baseline cardiovascular risk of the study participants using the Framingham Risk Score. The correct calculation increased the number of participants identified as having ≥15% 10-year risk from 5737 to 7089, a change from 61% to 76% of the total study population. This information is important for researchers attempting to validate and extend the trial’s findings and is particularly germane because the recently released American Heart Association/American College of Cardiology blood pressure guidelines changed blood pressure targets for pharmacologic therapy only for high-risk individuals.

## 1 Introduction

In April 2017, the *New England Journal of Medicine* (NEJM) hosted a summit on “Aligning Incentives for Sharing Clinical Trial Data,” with the aim of providing a demonstration of the benefits of clinical trial data sharing.^2^ In the months leading up to the summit, NEJM launched the SPRINT Data Analysis Challenge, which offered researchers access to the SPRINT clinical trial database in order to provide a real-world demonstration of the benefits of clinical trial data sharing. Investigators from around the world used the SPRINT data to produce research abstracts that were shared publicly. The SPRINT data are now available via the Biologic Specimen and Data Repository Information Coordinating Center (BioLINCC).^3^

SPRINT, a randomized clinical trial sponsored by the National Institutes of Health (NIH), compared a more intensive systolic blood pressure (BP) target ( <120 mm Hg) with a standard target ( <140 mm Hg) among non-diabetic patients aged 50 or older with hypertension, and with known cardiovascular disease or known elevated risk for cardiovascular disease.^2^ After observing significantly fewer cardiovascular events among patients allocated to the more intense treatment regimen, the Data Safety and Monitoring Board stopped the trial early, and the primary results were published in NEJM on November 26, 2015.

Our research group planned to use the SPRINT data. We first sought to determine if we could replicate the information in the main publication. As part of this effort, we calculated the 10-year Framingham Risk Scores (FRS). The published SPRINT paper reported that 61% of the participants were identified as having ≥15% 10-year risk based on the FRS.^1^ We found that this number did not match what we calculated from the data made available from BioLINCC. We emailed our findings to the coordinator of the NEJM Challenge from whom we received a response quoting a BioLINCC official, “The equation appears to have originally been calculated with the coefficients for treated systolic blood pressure and untreated systolic blood pressure reversed” (personal communication). In this report, we describe our mathematical analysis of this discrepancy and its implications.

## 2 Methods

### 2.1 Data Source

BioLINCC provided the data underlying the main publication of the SPRINT trial. These data were organized into in five data sets, including patient baseline information, blood pressure readings over time, primary and other outcomes, patient status at the end of intervention, and adverse events.

### 2.2 Data Variables

The variable of interest was the reported FRS, denoted by a variable labeled ‘*risk10yrs*’ in the baseline information data set. We also used the seven variables included in the FRS score: age, total cholesterol, HDL cholesterol, systolic blood pressure (SBP), antihypertensive medication use, current smoking status, and sex. As a result of the exclusion criterion, all SPRINT participants at baseline did not have diabetes, another FRS variable.

### 2.3 Risk Score Calculation

We calculated the FRS using the sex-specific formula derived originally from Cox proportional-hazards models in a 2008 paper by D’Agostino *et al.* using the 7 variables above (all of which were made available in the BioLINCC data).^4^ The NEJM Challenge coordinators confirmed that the D’Agostino *et al.* regression model (hereafter referred to as the “true” model) was appropriate for calculating the 10-year risk used in SPRINT (personal communication).

The continuous variables were (natural) log-transformed. Regression coefficients for each variable (shown in our Table 1) were calculated via the Cox model in the D’Agostino paper. If we represent the variables as *X_i_* (*X*_1_ is log(age), *X*_2_ is log(total cholesterol), etc.) and their corresponding coefficients as β*_i_*, then for each patient we can form the linear combination of the above variables and coefficients, given by Σβ*_i_X_i_*. With this calculated, the final risk score for women is given by

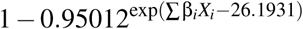

and for men by

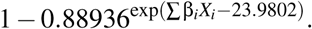

**Table 1.** Regression Coefficients for Cox regression model used to predict CVD risk^4^

| Variable | Women | Men |
| --- | --- | --- |
| Log of age | 2.32888 | 3.06117 |
| Log of total cholesterol | 1.20904 | 1.12370 |
| Log of HDL cholesterol | -0.70833 | -0.93263 |
| Log of SBP if not treated | 2.76157 | 1.93303 |
| Log of SBP if treated | 2.82263 | 1.99881 |
| Smoking | 0.52873 | 0.65451 |
| Diabetes | 0.69154 | 0.57367 |

### 2.4 Statistical Analysis

We compared our calculated FRS values with those in the *risk10yrs* variable. We then also calculated the percentage of participants with ≥15% 10-year risk by the calculated score and compared it with the ≥15% 10-year risk by the *risk10yrs* variable and the published result in the original SPRINT paper.^1^

To verify the validity of the explanation provided to us for the discrepancy, we tested whether interchanging the coefficients for treated SBP and untreated SBP resolved the discrepancy.

## 3 Results

We used the data from all 9361 study participants in SPRINT.

### 3.1 Comparison of *risk10yrs* with Published Result

Table 1 of the original SPRINT manuscript indicates that the number of participants whose FRS is ≥15% was 2870 and 2867 for intensive and standard treatment, respectively. Also, the mean ± standard deviation (SD) of the FRS values were 20.1 ± 10.9% and 20.1 ± 10.8% for intensive and standard treatment, respectively. All of these data agree with the numbers calculated using the *risk10yrs* variable provided to SPRINT Challenge participants.

### 3.2 Comparison of Calculated and Reported Risk Score

Our calculated FRS using the true model were not consistent with the reported scores in the *risk10yrs* variable. Specifically, 7089 (76%) of patients had ≥15% 10-year cardiovascular risk according to the calculated score versus 5737 (61%) using the score determined from the provided *risk10yrs* variable. The mean ± SD 10-year cardiovascular risk was 24.8 ± 12.5% for the calculated score versus 20.1 ± 10.9% for the score based on the *risk10yrs* variable.

The SPRINT Challenge variable *InclusionFRS*, derived from the *risk10yrs* variable, indicated that 5737 patients were included based on ≥15% 10-year risk. This number was consistent with the data presented in Table 1 of the original SPRINT manuscript, indicating a discrepancy between the results calculated from the SPRINT data and the published data in the SPRINT trial, which is consistent with the provided *risk10yrs* variable.^1,4^

We created a scatter plot (Figure 1) of the provided *risk10yrs* variable against our calculated FRS. We found our calculated FRS was lower than the *risk10yrs* variable for previously untreated patients and higher for previously treated patients. The overall effect, since 91% were previously treated and treatment is associated with higher risk, was to represent the study sample as being lower risk than was true.

**Figure 1:**
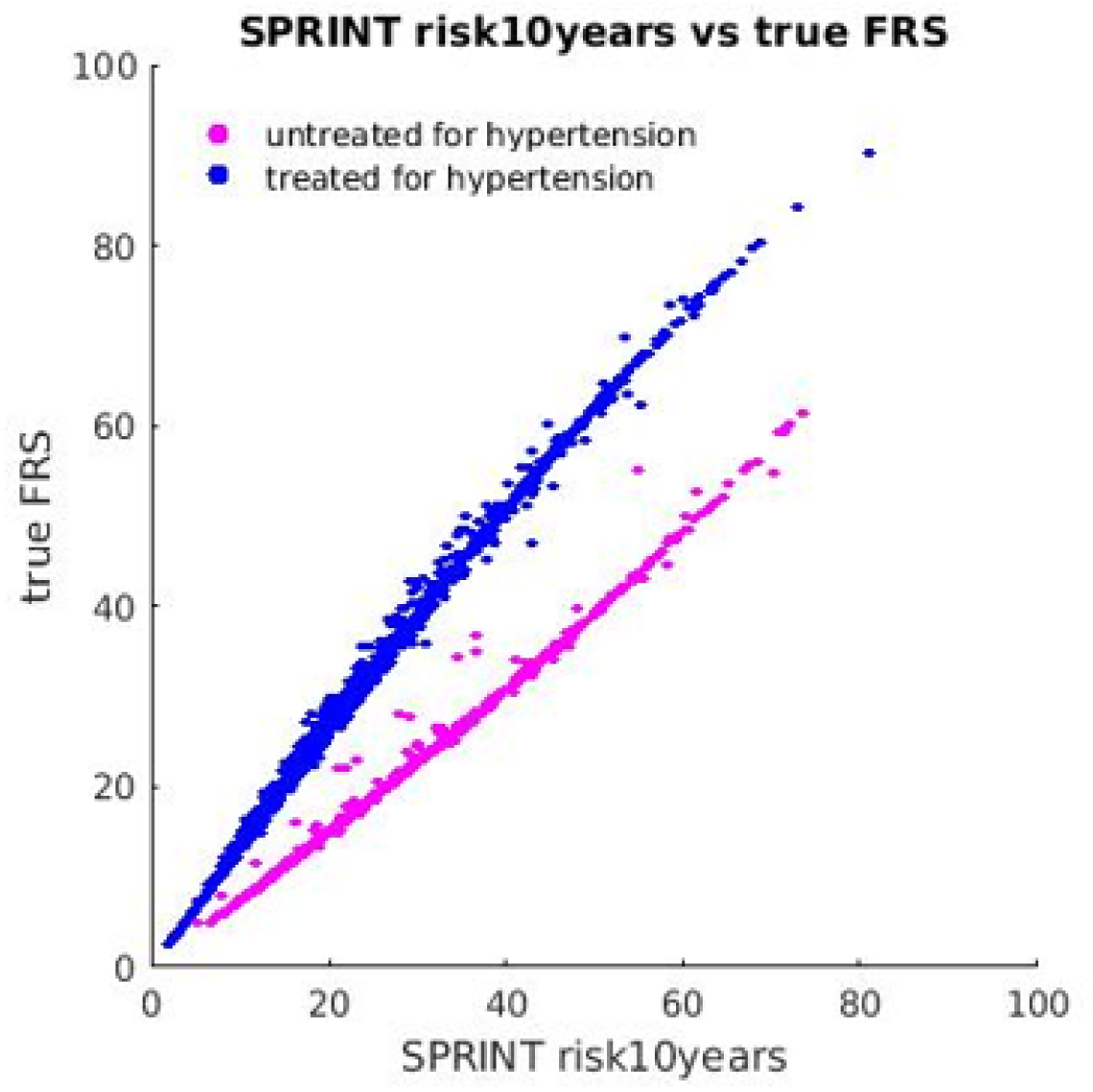
Scatter plot of the *risk10years* variable vs. the calculated variable using the true model^4^. Color is used to indicate the correct blood pressure treatment status of study participants. This figure illustrates the effect of interchanging the prior antihypertensive medication use variable – the FRS is under-estimated for those being treated and overestimated for the untreated population.

To verify the effect reversing the coefficient for treated systolic blood pressure and untreated systolic blood pressure in the FRS true calculation (to test if this accounts for the discrepancy), we tested the effect of reversing the coefficients for treated systolic blood pressure and untreated systolic blood pressure formula in our calculated FRS. After doing so, there remained 585 (6.3%) participants for whom the FRS results from the published paper differed from those calculated using SPRINT data with reversed treatment coefficients. For 10 of these participants, the risk reported in *risk10yrs* agrees exactly with the correct formula and not the formula with the reversed coefficients. For the remaining 575 of these participants, we are unable to either replicate or explain the published FRS results.

## 4 Discussion

After receiving access to the data underlying SPRINT through the SPRINT Data Analysis Challenge, we found an error in the FRS calculations in the SPRINT publication.^1^ The SPRINT main paper erroneously stated that 61% of patients had ≥15% 10-year cardiovascular risk, instead of the true value of 76%. The new ACC/AHA Blood Pressure Guidelines relied heavily on SPRINT in making recommendations to lower the treatment target for which pharmacologic therapy should be initiated for highrisk individuals.^5^ Of note, this information does not change the results of the trial, but actually shows that the study is particularly relevant to high-risk individuals since even more were higher risk according to the FRS. The finding also provides support to the decision by the National Institutes of Health and the SPRINT investigators to share their data by showing a benefit of data sharing. The error in the main paper has now been corrected.^1^

The FRS was one of the four eligibility criteria for SPRINT, and the most common (accounting for eligibility of 61.3% of all study participants). Therefore, understanding the subset of study participants at ≥15% risk at enrollment is critical to understanding the SPRINT study population and considering the real-world population to whom the study results can be generalized.

Some questions persist. BioLINCC has stated that the population at ≥15% risk was determined at a pre-baseline screening visit (which was not reported in the main paper), but these data were not available to SPRINT Challenge participants.^6^ Therefore, it is not possible to determine the effect of the incorrect calculation on the inclusion criteria and whether it was used to make these determinations. Additionally, the reversing of the coefficients for treatment, which was suggested as the coding error responsible for the error, does not fully explain the discrepancy.

This correction highlights an often-overlooked benefit of data sharing in medicine: error identification and correction by reproducing research to verify previously published research findings. Many researchers report having failed to reproduce their own scientific experiments, or an experiment of a colleague, and are just now beginning to establish procedures to foster scientific reproducibility.^7^ Clinical trial data sharing is likely the best method to facilitate reproducibility in the clinical sciences. The sharing of data can enable the wisdom of crowds to emerge, proper questioning and clarification of methods and, ultimately a greater understanding of particular studies. Moreover, sharing empowers other researchers to ensure that the contributions of the patient participants and scientists who create a study are honored by generating as much clinically -and scientifically-relevant knowledge from the study as possible.

In conclusion, the SPRINT Data Analysis Challenge demonstrated how clinical trial data sharing enables increased knowledge generation to improve clinical practice and scientific understanding. Our correction illustrates a secondary benefit to data sharing, namely that data sharing allows for outside researchers to reproduce existing analyses, and in that process, discover any errors. Even in this highly curated, limited dataset, known to be shared with the public and constructed by experts in the field, we found an error that was likely the result of a simple miscode for most patients. Greater availability of clinical trial protocols and underlying datasets will allow for novel investigations as well as greater verification and reproducibility of existing investigations, strengthening confidence in trial results and conclusions.

